# Computational modelling of novelty detection in the mismatch negativity protocols and its impairments in schizophrenia

**DOI:** 10.1101/2025.09.30.679453

**Authors:** Ahmed Eissa, Jan Fredrik Kismul, Atle Bråthen Pentz, Torbjorn Elvsåshagen, Christoph Metzner, Ibrahim Akkouh, Srdjan Djurovic, Alexey Shadrin, Marja-Leena Linne, Gaute T. Einevoll, Ole A. Andreassen, Tuomo Mäki-Marttunen

## Abstract

The human auditory system rapidly distinguishes between novel and familiar sounds, a process reflected in mismatch negativity (MMN), an EEG-based biomarker of auditory novelty detection. MMN is impaired in psychiatric conditions, most notably schizophrenia (SCZ), yet the neuronal mechanisms underlying this deficit remain unclear. Here, we combined computational modelling and genetic analyses to investigate how SCZ-associated cellular abnormalities affect auditory novelty detection. We developed an integrate-and-fire spiking network model capable of detecting four types of auditory novelty, including stimulus omission. Based on assumptions of short-term depressing synapses between the subpopulations of the network and the existence of neuronal inputs that are phase-locked to the rhythm of the recently experienced stimulus sequence, the model reliably reproduced MMN-like novelty detection and allowed systematic testing of SCZ-related cellular alterations. Simulations revealed that both reduced pyramidal cell excitability, linked to ion-channel dysfunction, and decreased spine density impaired novelty detection, with the latter producing stronger deficits. Our work provides a flexible spiking network model of auditory novelty detection that can link cellular-level abnormalities to measurable MMN deficits, improving their mechanistic interpretation and helping to explain the heterogeneity of SCZ.

## 1 Introduction

The human auditory system is highly effective in distinguishing between novel and familiar sounds and auditory patterns. Detecting novel auditory stimulus can help us shift attention and plan our actions accordingly, but it also generates eventrelated potentials (ERPs) that can be measured using an electroencephalogram (EEG) in both conscious and unconscious modes of operation [Alho, 1992]. Mismatch negativity (MMN), defined as the difference between ERPs elicited by novel and familiar tones, is the most widely EEG-based biomarker of auditory novelty detection [Näätänen, 1982, Lakatos et al., 2020]. MMN is clinically relevant because it is impaired in psychiatric disorders, particularly in schizophrenia (SCZ) [Light and Näätänen, 2013]. However, due to the complexity of the auditory system, the neuronal mechanisms of auditory novelty detection in general, as well as its impairment in SCZ, remain incompletely understood. Elucidating these mechanisms may help bridge cellular-level alterations associated with SCZ to clinically observed deficits.

In this work, we employed computational modelling of spiking neuronal networks to investigate the auditory novelty detection and the ways in which it is compromised by SCZ-associated cellular-level abnormalities. We developed an integrate-and-fire (IAF) model network capable of detecting four types of auditory novelty frequently used in the MMN paradigm: a frequency deviant, an omitted stimulus, a deviant with a longer duration than the standard tone (henceforth the “duration deviant”), and, finally, a deviant with a shorter duration than the standard tone (henceforth the “inverse duration deviant”). We demonstrated the robustness of our proposed network model in novelty detection under a variety of conditions and examined how SCZ-associated reductions in pyramidal cell excitability, suggested by alterations in ion-channel expression [Hoffman et al., 2019, Mäki-Marttunen et al., 2024], and a decreases in spine density [Shelton et al., 2015] affect novelty detection. Although both alterations impaired deviance detection, our simulations suggested that decreased spine density exerts stronger effects than decreased neuronal excitability. This work therefore serves a dual purpose: to introduce a mechanistic spiking network framework for auditory novelty detection, and to apply it in the context of SCZ to identify how cellular abnormalities may translate into measurable MMN deficits. By linking cellular-level alterations to measurable impairments in auditory novelty detection, our model provides a framework for developing targeted therapeutic strategies and stratifying patients based on distinct neurophysiological profiles.

## 2 Methods

### 2.1 Network model for novelty detection in the auditory pathway

The MMN signal can be registered using many different protocols, where the deviant sound is different from the standard sound in terms of pitch, phoneme (if human voice-based stimuli used), duration, inter-stimulus interval, or intensity, or even when the stimulus is omitted [Garrido et al., 2009]. The aim of this work was to present a spiking neuronal network model capable of detecting mismatches of four types, namely, a frequency deviant, an omitted stimulus, a duration deviant, and an inverse duration deviant. To describe the neuronal mechanisms that permit novelty detection, we introduced a spiking neuronal network of multiple neuron populations.

We made the following assumptions (their validity is discussed in the Discussion):

1. There are excitatory and inhibitory populations of neurons that selectively respond to each tone, and the excitatory neuron populations project to other populations by short-term depressing synapses.
2. Some of these populations respond upon receiving either short or long-lived auditory stimulus, while some populations require a long-lived auditory stimulus to be activated.
3. There are populations of neurons (separate from the above) that are phase-locked to a 2-Hz stimulus presentation frequency with different phase shifts.
4. These neuron populations interact with an “output” population that detects auditory mismatches of different type.

To develop a network model to meet this aim under the above assumptions, we considered excitatory (ES) and inhibitory (IS) populations that selectively responded to the standard (S) tone, and excitatory (ED) and inhibitory (ID) populations that selectively responded to the deviant (D) tone. We also modelled two delayed-activating populations (ESD, EDD) that were similar to ES and ED but required longer inputs to fire. To activate these neurons, we used a stimulus that lasted twice as long as the standard tone and ended at the same time. Furthermore, we modelled an excitatory neuron population that received inputs from a population that was phase-locked to a 2-Hz stimulus presentation rate with a phase that was coincident with standard stimulus (EP), as well as another population (EP2) with a phase aligned with the first half of the longer-duration deviant. Finally, we modelled an excitatory population that received inputs from all six excitatory populations and functions as an output population (EO). All excitatory synaptic connections were assumed to be short-term depressing, which allowed the repeated standards to induce a different network response than the deviants. The network is illustrated in Fig. 1A; step-by-step schematics are shown in Figs. S1–S4.

**Figure 1.**
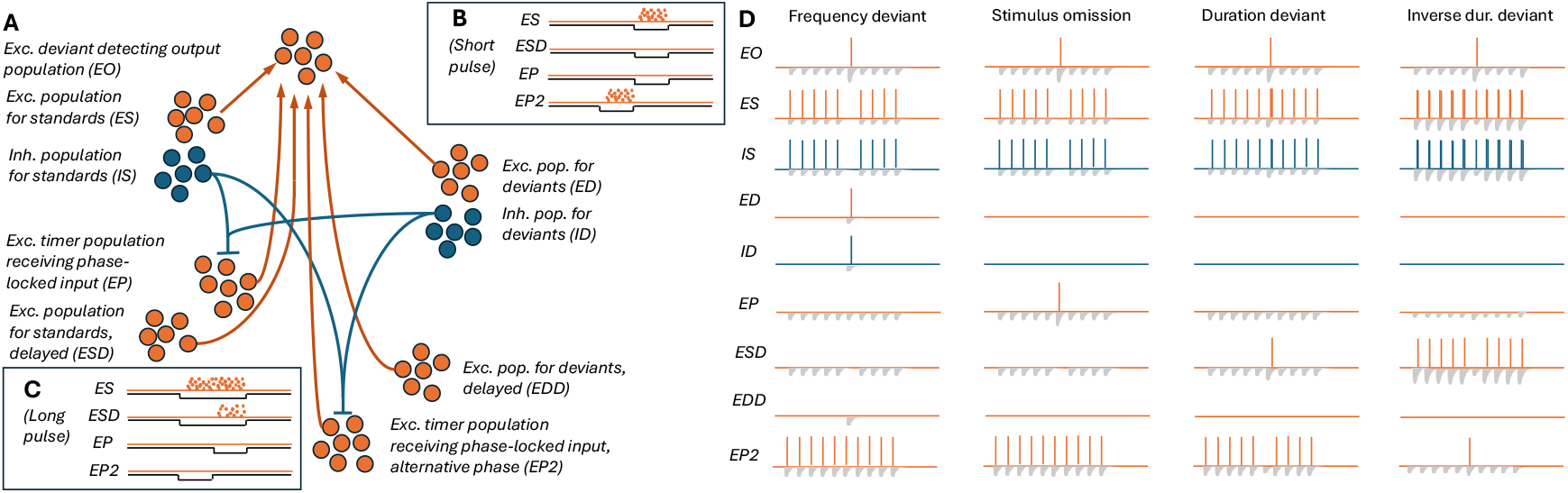
Illustration of the model network and the hypothesized mechanisms of deviance detection. See Fig. S1–S4 for a breakdown of the hypothesized mechanisms.

We here modelled only one additional specific frequency-tuned population in addition to the standard tone-tuned population and a pacemaking population tuned to particular phases of a particular stimulus presentation rate (2 Hz), but our framework can be extended to describe any number of such deviants and pacemaking signals.

We hypothesized that the network described above could perform the deviance detection by the following mechanisms. First, the ES and ED populations project to the output EO population via depressing synapses. Repeated standards elicit weaker responses than deviants, such that EO fires to a deviant but not to repeated standards (Supp. Fig. S1). Second, the omission detecting population EP receives excitatory inputs from phase-locked neurons (not explicitly modelled) that were assumed to be independent of the auditory stimuli (however, the activation of these neurons may be dependent on the history of the auditory stimuli through plasticity mechanisms). These inputs are counteracted by inhibitory inputs from the frequency-sensitive neuron populations IS and ID, and thus, the EP population only fires when none of the frequency-sensitive neuron populations are active (Fig. 1B). When the omission detecting population EP fires, it activates the output population (Supp. Fig. S2). Third, the delayed-activating population ESD fires in response to a long but not short stimulus (Fig. 1B–C). At the introduction of the first longer stimulus among shorter standards, the ESD fires and makes the output population fire as well (Supp. Fig. S3). Fourth, the long stimulus inhibits not only the EP but also the EP2 population (Fig. 1C) that would otherwise fire at the phase preceding the phase of the EP population (Fig. 1B). Similar to EP, the EP2 population projects to the output population with short-term depressing synapses. EP2 thus remains silent in response to recurring long standards but fires when a shorter duration stimulus is used as a deviant (Fig. 1C) and causes the output population to fire (Fig. S4). These mechanisms are illustrated in Fig. 1D — see Results for our approach for fitting the model to reproduce these mechanisms.

We modelled the neurons as leaky IAF units that received AMPAR- and NMDAR-mediated and GABAergic conductance-based synaptic inputs in addition to square-pulse currents that were timed to correspond to inputs from stimulus-frequency-coding and pacemaker neurons. Each population consisted of N=40 neurons. The glutamatergic inputs from the ES, ED, EP, ESD, EDD, and EP2 populations to the output population EO were short-term depressing according to the following scheme (adapted from [Wang, 1999]):

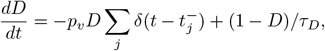

where *D* is the fraction of releasable vesicles, *p*_*v*_ is the release fraction per spike, and *τ*_*D*_ is the time constant of recovery from depression. The term 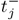 represents the time instants of the presynaptic spikes.

### 2.2 Optimization of the model parameters

We used the grid search method to achieve model parameters that yielded a good deviance detection performance. We ran the four MMN experiments (the frequency deviant, the omission of stimulus, the duration deviant, and the inverse duration deviant) for a large number of parameter sets (5 stimulus amplitudes × 5 exc. synaptic conductances from the ES and ED populations × 5 exc. synaptic conductances from the EP and EP2 populations × 7 exc. synaptic conductances from the ESD and EDD populations × 2 NMDA/AMPA ratios × 6 inh. synaptic conductances × 2 depression strengths × 2 membrane capacitances for the ESD and EDD populations; 42000 parameter sets in total). The tested parameter ranges are shown in Table S2A). We considered networks where ≥80% of the output (EO) neurons fired once or more for the deviant and the number of EO spikes following the deviant was at least 6 times an average number of EO spikes following a standard as acceptable models.

### 2.3 Quantification of the deviance detection

To assess the network activity, we quantified deviance detection using an index inspired by the MMN, calculating the average numbers of spikes induced by the deviant stimulus (or omission of stimulus) (*N*_*deviant*_) and that induced by the standard (*N*_*standard*_) in the output population. For each stimulus, we included the spikes elicited from 150 ms prior to the expected onset of the stimulus to 350 ms after the onset of the stimulus and obtained the firing rates *f*_*deviant*_ and *f*_*standard*_ by normalising the numbers of spikes by the interval (0.5 sec). We then defined the deviance detection index as

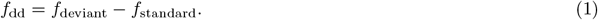

#### 2.4 Modelling of spike-timing-dependent plasticity (STDP)

In Section 3.3, we tested the entrainment of a neuron receiving inputs from many different phase-locked populations to external auditory stimuli arriving at a certain rhythm and phase via STDP. We implemented the synapse weight update rules using the standard STDP formalism available in Brian2. For this experiment, we used an IAF neuron with a normalized membrane potential (resting membrane potential at 0 and threshold potential at 1 A.U.). The state of each synapse was characterized by dynamic variables representing presynaptic and postsynaptic traces, which decayed exponentially with distinct time constants (20 ms for pre-post and 40 ms for post-pre interactions). When a presynaptic spike occurred, the postsynaptic membrane potential was incremented proportionally to the current synaptic weight, the presynaptic trace was increased, and the weight was potentiated depending on the postsynaptic trace. Conversely, when the postsynaptic neuron fired, its trace was incremented and synaptic weights were depressed depending on the presynaptic trace. Weight updates were clipped to remain within fixed bounds (0 to 1). The learning rate parameters were set asymmetrically (*A*_plus_ = 0.015, *A*_minus_ = -0.045) to reflect a biologically realistic bias toward depression, and initial synaptic weights were set to 0.05. This implementation allowed synaptic efficacy to evolve according to the relative timing of pre- and postsynaptic activity, following a standard pair-based STDP rule.

### 2.5 Use of genetic data in the modelling

To study the effects of altered expression of ion channels on deviance detection, we used the CommonMind RNA expression data [Hoffman et al., 2019] from two regions: the anterior cingulate cortex (ACC, N=478 of which 251 were controls and 227 SCZ patients) and prefrontal cortex (PFC, N=426 of which 215 were controls and 211 SCZ patients). We followed the approach of [Mäki-Marttunen et al., 2024] in applying these data for modelling. Unlike the study of [Mäki-Marttunen et al., 2024] that employed biochemically and biophysically detailed neuron modelling, here we used simplified IAF models that do not permit information on ion channel expression to be integrated into the model. We thus used the Hay model [Hay et al., 2011] as a proxy to estimate the effects of altered ion channel expression on pyramidal cell firing behaviour.

In short, we first used a list of all SCZ risk genes associated with fast or slow neurotransmission [Devor et al., 2017] and the sister genes in their gene families (Table S2 in [Mäki-Marttunen et al., 2024]). We then extracted the high-risk variants within these genes from [Trubetskoy et al., 2022]. While the study of [Trubetskoy et al., 2022] applied several complementary strategies to assign SNPs to credible genes, in our analysis we used a simpler positional approach: SNPs were assigned to genes if their coordinates fell within gene boundaries. Genes were retained as genes of interest if at least one SNP within their boundaries had a p-value smaller than 5 · 10^*−*6^. In particular, we focused on the genes whose effects we could model using the model of [Hay et al., 2011] (Table S1A). We complemented this set of genes by sets of genes that were differentially expressed in SCZ compared to healthy controls (HC) (Table S1B). To do this, we analysed the CommonMind data of post-mortem RNA expression in PFC and ACC. The expression data were normalized using the DESeq2 method [Love et al., 2014]. We also employed an imputation tool, CIBERSORTx [Steen et al., 2020], to obtain estimates of neuronal expression in these brain regions instead of the bulk expression using a reference single-cell RNA dataset [Zhang et al., 2016]. We only considered the genes that were successfully imputed, and furthermore, we restricted our analysis on genes that showed a “medium” or “high” level of protein expression in neuronal cells of cerebral cortex or had a high specificity for expression in excitatory neurons according to the single-cell RNA sequencing data of Human Protein Atlas. We then normalised each gene expression of each subject by the average expression of the underlying gene in the 251 (ACC) or 215 (PFC) controls. When there were multiple SCZ hit genes affecting a single model parameter, we used the average of the expression-level factors to obtain a single factor.

The pyramidal cell model of [Hay et al., 2011] consisted of 196 compartments and described the following ionic currents: high-voltage activated Ca^2+^ current (*I*_Ca,HVA_), low-voltage activated (LVA) Ca^2+^ current (*I*_Ca,LVA_), non-specific cationic current (*I*_*h*_), transient (*I*_Na,t_) and persistent (*I*_Na,p_) Na^+^ currents, muscarinic K^+^ current (*I*_*m*_), transient (*I*_K,t_) and persistent (*I*_K,p_) K^+^ currents, Ca^2+^-activated SK current (*I*_SK_), Kv3.1-mediated K^+^ current *I*_Kv3.1_, and the leak current (*I*_leak_). Changes in the expression of the subunits contributing to these currents were modelled by altered maximal conductance 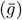 of the underlying current. Similar to [Mäki-Marttunen et al., 2024], we attributed the expression of CACNA1C and CACNA1D to *I*_Ca,HVA_, that of CACNA1I to *I*_Ca,LVA_, that of HCN1 to *I*_*h*_, and that of KCNQ3 to *I*_*m*_, those of KCNB1 and KCND3 to *I*_K,p_, and that of SCN1B to *I*_Na,t_.

To determine the effects of altered expression on intrinsic excitability, we simulated the pyramidal cell spiking response to a set of somatic currents (0 to 1.0 nA in intervals of 0.1 nA). We simulated the model separately for each subject, where the conductances were adapted in a subject- and current species-specific manner as explained above. We determined the f-I curves, i.e., the spiking frequencies during the last 15.5 sec of the current injection with respect to the current amplitude. The obtained results, and in particular, the obtained subject-wise areas under curve (AUC_*i*_), were then refined for further use in IAF simulations in one of the two ways:

1. The membrane capacitance of the IAF model for subject *i* was multiplied by 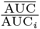 2, where 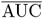 is the average AUC across control subjects.
2. The membrane capacitance of the cortex-adapted IAF model for subject *i* was fitted to yield the exactly same AUC_*i*_ in a corresponding f-I experiment with the IAF model.

We then averaged across the membrane capacitances of the SCZ population to yield an estimate of how much the SCZ diagnosis, mediated by altered ion channel expression, affects the membrane capacitance.

### 2.6 Code availability

All simulations were run using NEURON v. 8.2.6, using Python (3.9.20) interface. Our simulation scripts are available at ModelDB, accession number 2019882 (https://modeldb.science/2019882, password ‘mmn’ required during peer review).

## 3 Results

### 3.1 Development of a spiking network model capable of novelty detection in the frequency and omission-based MMN

To describe the novelty detection based on spiking neuronal network dynamics in health and disease, we introduced a network of multiple neuron populations (see Methods and Fig. 1). We first ran a series of grid search experiments to find an approximate range of parameters (stimulus amplitudes, synaptic conductances, membrane capacitances, and the depression strength) within which appropriate spiking and a different response for deviants versus standards was achieved. We then ran a large grid search to optimise the novelty detection (see Table S2A). We achieved strong deviance detection for 7249, 10118, 15022, and 7199 parameter sets (out of 42000) in the frequency-deviant, stimulus-omission, duration deviant, and the inverse duration deviant protocols, respectively (Fig. 2A). 16 parameter sets supported detection across all four protocols (Fig. 2A) — these parameters are listed in Table S2B. The behaviour of all nine neuron populations in response to the four MMN protocols is illustrated in Fig. 2B.

**Figure 2.**
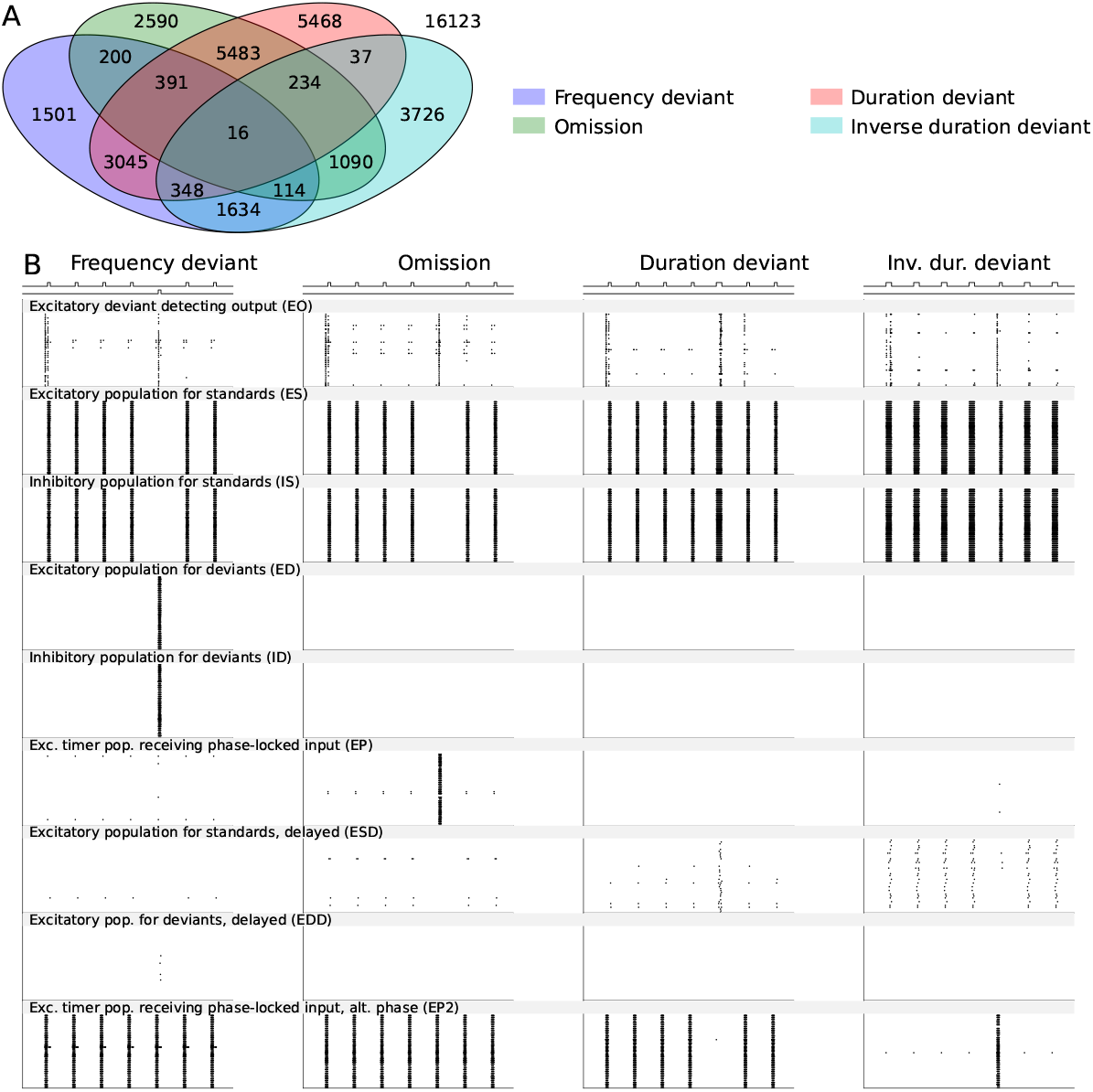
Fitting of the spiking network model for novelty detection in four different types of MMN protocols and the firing behaviour of the subpopulations in the four protocols. **A**: The Venn diagram showing the number of parameter sets (out of 42’000) that yielded an acceptable novelty detection in the frequency deviant (blue), omission (green), duration deviant (red), and inverse duration deviant (cyan) MMN protocols and their intersection. 16 parameter sets yielded an acceptable novelty detection in all four protocols. **B**: Population spike raster plots of the nine subpopulations (40 neurons in each) in the four MMN protocols according to one of the 16 acceptable models.

Taken together, our grid-search method yielded spiking networks that effectively detected novelty in typical MMN stimulus protocols.

### 3.2 Robustness

To show the robustness of the novelty detecting network, we validated the model by additional experiments. First, we showed that the network can detect novelty in a roving deviant paradigm where a sequence of five identical stimuli (whether short or long tones of the standard or deviant frequency) forms a new standard after each transition (Fig. 3A–B). In the same experiment, we also tested the omission of stimulus when each of the four types of tones were used as a standard (Fig. 3A–B). The deviance detection indices were of similar magnitude or larger in these transitions compared to the MMN protocols used in Fig. 2 (Fig. 3C).

**Figure 3.**
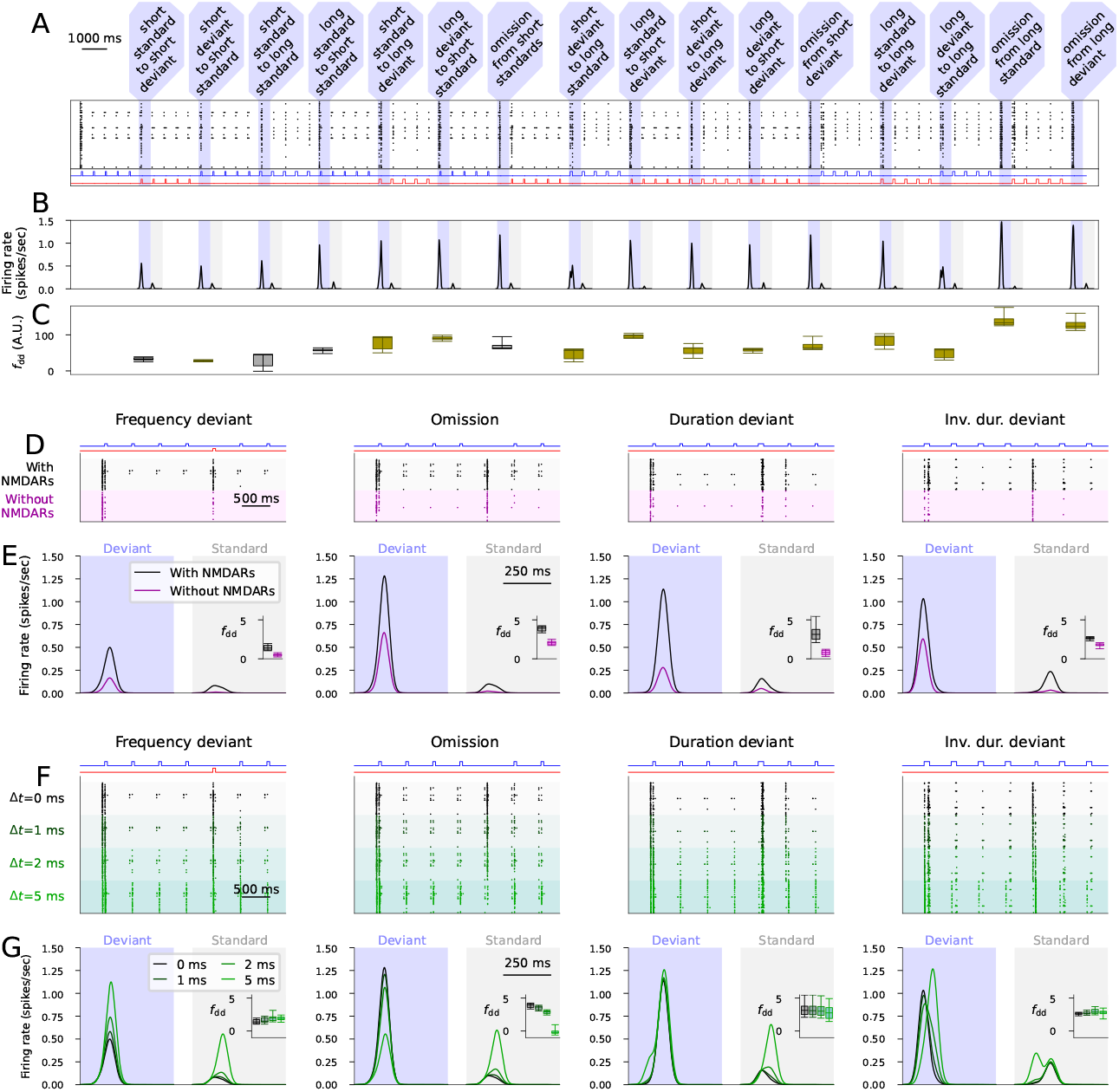
The novelty detecting network functions well in the roving paradigm, shows NMDAR-dependency, and is robust against introduction of synaptic delays. **A–C:**Experiments with an alternative stimulation sequence where the four stimuli (short standard, short deviant, long standard, long deviant) are repeated in sequences of five tones and all possible transitions between them and all omissions are included. **A**: The output population activity according to the model with the default parameter set. The shaded areas represent the deviants, i.e., transitions to a new type of tone or omissions. The insets below the graph show the stimuli corresponding to the standard (blue) and deviant (red) tones across time. **B**: The experiment of (A) was repeated for all 16 models. The plots show the average firing rate curves induced by the deviants (darker gray) and those induced by the previous tone (light gray). **C**: Box plots of the deviance detection index determined from the data of (B). The magenta bars represent the MMN protocols used in Fig. 2, and the cyan bars represent the alternative deviants not used for fitting the model. **D–E**: The deviance detection with and without NMDAR-mediated currents. **D**: The output population activity according to the model with the default parameter set in the presence (black) and absence (magenta) of NMDAR-mediated currents. **E**: Average (across the 16 models and 10 repetitions with different random number seeds) firing rate curves induced by the deviants (darker gray) and those induced by the previous tone (light gray) in the four MMN protocols in the presence (black) and absence (magenta) of NMDAR-mediated currents. The insets show the box plots of the deviance detection indices in each protocol. **F–G**: The deviance detection with and without synaptic delays. **F**: The output population activity according to the model with the default parameter set in the absence (black) and presence of 1 (dark green), 2 (green) or 5 (light green) ms synaptic delays. **G**: Average (across the 16 models and 10 repetitions with different random number seeds) firing rate curves in the four MMN protocols in the absence (black) and presence (green) of synaptic delays. The insets show the box plots of the deviance detection indices in each protocol.

We also simulated the deviance detection in all four MMN protocols when the NMDARs were blocked. Similar to experimental observations [Farley et al., 2010], the simulations with NMDAR blockage showed a decreased response to both deviants and standards (Fig. 3D–E). Our simulations, however, suggested that the deviance detection index was also decreased in the absence of NMDAR activation (Fig. 3E).

Biological networks include delays consisting of axonal and dendritic signal conduction and a presynaptic neurotransmitter release before a presynaptic AP induces AMPAR, NMDAR or GABAR-mediated currents in the postsynaptic neuron, which is not taken into account in the model. We tested the functionality of the novelty detection network in presence of neurotransmission delays. The deviance detection remains unchanged in all MMN protocols when small delays (Δ*t*=1 or 2 ms) were used, but the detection of stimulus omission was lost when a large delay (Δ*t*=5 ms) was used (Fig. 3F–G).

We next tested the contribution of individual populations in the deviance detection. In these experiments, we removed some of the populations and performed the four different experiments without the selected populations and recorded the output population firing activity. When the EP2 population that was responsive during the alternative phase of the 2- Hz stimulus presentation was removed, the detection of inverse duration deviants was lost while the other three types of deviants were detected (Fig. S5A). Removing both EP and EP2 abolished omission and inverse duration detection but preserved frequency and duration detection (Fig. S5B). By contrast, removing the delayed-activating populations (ESD, EDD) alongside the EP2 population abolished the detection of all duration deviants but preserved the detection of frequency deviants and omitted stimuli (Fig. S5C). When only the tone-sensitive populations (ES, ED, IS, and ID) and the output population were in place while others were removed, the network only detected the frequency deviants (Fig. S5D).

We verified that the network was robust against small changes in network size and variability in neuronal excitability (Fig. S6A–B). The network detected deviants when its size was halved (20 neurons) or increased to 60 neurons (Fig. S6A), and when the variability of the membrane capacitance of the neurons was reduced to 0.2× mean(*C*_*m*_) or increased to 0.4× mean(*C*_*m*_) (Fig. S6B). However, given too little variability in the membrane capacitance of the neurons (SD(*τ*) ≤ 0.1× mean(*τ*)), the inverse duration deviants were no more detected (Fig. S6B). We also made sure the network does not produce a deviance detection artifact due to the phase-locked neuron activity when random auditory stimuli were given as inputs instead of the ryhthmic tone inputs (Fig. S6C).

While the activation of the stimulus-encoding neuron populations (ES, ED) can be expected to be timely, the phase-locked populations may show large deviation in the timing of its activity. Here, we show that the detection of omitted stimuli is not dependent on a perfectly timed activation of the 2-Hz phase-locked populations. To do this, we randomized a phase for each of the phase-locked neuron from a uniform distribution [*t*_0_ − *θ, t*_0_ + *θ*] where the jitter magnitude term *θ* ranged from 0 (no jitter) to 20 ms. A jitter of ±2 to 5 ms had little effect on the deviance detection in any of the MMN protocols, but a jitter of ±20 ms almost abolished the detection of omissions and also mildly impaired the detection of inverse duration deviants (Fig. S6D–E).

Taken together, our network model for deviance detection behaves in a predicted manner in a roving paradigm and when certain populations are removed. Although our network is simplified in many aspects, the network functions robustly when uncertainties that typically appear in biology, such as delays and parameter variabilities, are added.

### 3.3 Phase-locking to the stimuli in the MMN protocol can arise from STDP and rhythmically active neurons

The neurons phase-locked to the stimulus frequency were required in our network model for the detection of omissions, but it is unlikely that the auditory pathway constantly has neurons that are phase-locked to all possible frequencies and multiple phases of these rhythms with a high precision. Here, we show that phase locking to rhythmic entrainment stimulus can emerge as a result of STDP in the presence of rhythmically active neurons.

We simulated a network consisting of 164 neurons that were rhythmically active with 41 different frequencies (1 to 3 Hz in steps of 0.05 Hz) and ten different phases (0°, 36°, …, 324°) and a postsynaptic neuron that received inputs subject to STDP from these neurons (Fig. 4A). At baseline, the APs in the 164 presynaptic neurons failed to induce APs in the postsynaptic neuron (Fig. 4B). We then made the postsynaptic neuron fire with strong, rhythmic entrainment stimuli at a rate of 2 Hz. During the entrainment, the synaptic connections from the neurons that fired after the AP in the postsynaptic neuron were weakened, while the connections from neurons that most often fired before the postsynaptic AP increased in strength (Fig. 4C). This resulted in strong connections from a few neurons that were able to keep the postsynaptic neuron firing at the timed intervals even after the cessation of the entrainment stimulus (Fig. 4B). The strengthening of the synapse belonging to the optimally timed neuron was relatively large also when a small percentage of the entrainment stimuli was omitted similar to the MMN protocols (Fig. 4D–H): the synapse weight was 0.86 A.U. in the standard-only protocol (Fig. 4C–D) and 0.83 (Fig. 4E), 0.76 (Fig. 4F), or 0.72 A.U. (Fig. 4G) when every 20th, 10th, or 6th stimulus was omitted, respectively. If every fourth or second stimulus was omitted, the optimally timed synapse weight stabilized at 0.64 (Fig. 4H) or 0.48 (Fig. 4I), respectively.

**Figure 4.**
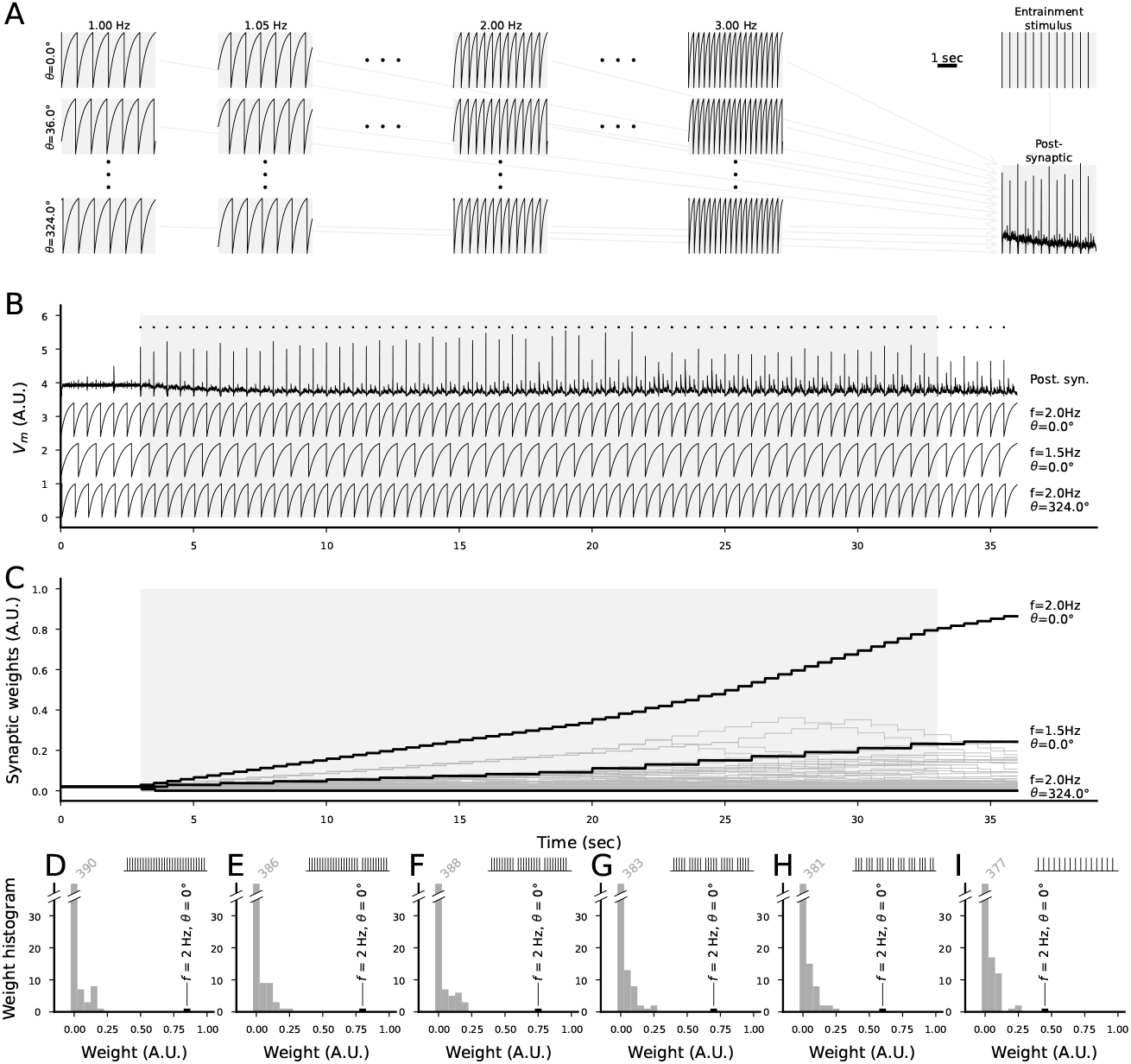
STDP-exhibiting synapses from rhythmically active neurons to a postsynaptic neuron receiving rhythmic entrainment stimulus can strengthen the synapses from the presynaptic neuron firing at the corresponding rate and optimal phase. **A**: Illustration of the network model. The model consists of 41 × 10 presynaptic neurons that are rhythmically active (and do not receive any inputs) with 41 different rates (1 to 3 Hz in steps of 0.05 Hz) and ten equally spaced phases, and a postsynaptic neuron that receives (initially) weak synaptic inputs from each presynaptic neuron and a strong entrainment input (e.g., caused by standard stimuli in the auditory MMN protocol). The subpanels show the membrane potential time courses of the postsynaptic neuron (bottom right) and 12 selected presynaptic neurons (left). **B**: Membrane potential time courses of the postsynaptic neuron (top), optimally and suboptimally phased presynaptic neuron (2nd and 3rd from top; the neurons whose synaptic strength grew most and second most during the entrainment), and the neuron whose synaptic strength decreased the most (bottom). The APs of the postsynaptic neuron are marked with dots above the membrane potential time course. The gray area depicts the time of the entrainment stimulus. The postsynaptic neuron does not fire APs before the entrainment stimuli, but fires rhythmically both during and after the entrainment. **C**: Time courses of the synaptic weights of all 410 synapses (dim). The synapses from the optimally and suboptimally phased presynaptic neuron and the neuron whose synaptic strength decreased the most (same as in (B)) are plotted in thick black lines. **D–I**: Histograms of the post-entrainment synaptic weights of all 410 synapses when none of the stimuli (D) or every 20th (E), 10th (F), 6th (G), 4th (H) or 2nd (I) entrainment stimulus was omitted. The black bar indicates the synapse from the optimally phased presynaptic neuron (2 Hz, fires at phase 0°, i.e., always 5 ms before the entrainment stimulus). The histograms are clipped at 40 synapses — the numbers of synapses in the smallest bin (weight between 0 and 0.05) are indicated above the bar.

Taken together, our analysis with rhythmically active point neurons connected with STDP-exhibiting synapses to a postsynaptic neuron receiving an entrainment stimulus suggests that the synapses from the neurons that fire in an optimally timed manner can be significantly strengthened due to entrainment (by, e.g., standard auditory stimuli in the MMN protocol) while other synapses are weakened. This synaptic strengthening may have the effect of making the postsynaptic neuron fire with the rate and phase of the entrainment stimulus even after the cessation of the stimuli, and it is not significantly impaired by omitting a small percentage of the entrainment stimuli.

### 3.4 Effects of SCZ cellular level phenotypes on novelty detection: the cortical detection hypothesis

The network model developed in Section 3.1 is abstract and not directly matched to a specific brain area. Although detection of novelty in auditory stimuli and generation of MMN is likely to be dependent on many brain areas across the whole brain, previous works have highlighted either the contribution of either cortical [Näätänen et al., 2007, Fishman and Steinschneider, 2012] or subcortical [Pérez-González et al., 2005, Malmierca et al., 2009] areas. Since subcortical neurons and circuits are different from their cortical counterparts by both structure and function, the way novelty detection is altered in the schizophrenic brain should be crucially affected by whether the novelty detection takes place in the cortex or subcortical regions. In this Section, we explore the effects of SCZ-associated cellular-level phenotypes on the predicted novelty detection under the assumption that *all modelled neuron populations are cortical neurons*.

First, we used the CommonMind data [Hoffman et al., 2019] that included the bulk RNA expression of ion-channel-encoding genes in the prefrontal cortices (PFC) of 426 subjects (211 SCZ, 215 HC) and anterior cingulate cortices (ACC) of 478 subjects (227 SCZ, 251 HC). We normalized the expression levels by DESeq2 [Love et al., 2014] and used CIBERSORTx [Steen et al., 2020] to obtain estimates of neuronal expression in these brain regions with the help of a reference singlecell RNA dataset [Zhang et al., 2016]. We focused on genes implicated by GWAS [Trubetskoy et al., 2022] or differential expression that were likely to be expressed in the cortex as done previously [Mäki-Marttunen et al., 2024] — see Table S1. We calculated subject-wise factors for each parameter of the Hay model [Hay et al., 2011] affected by these genes. We then performed the subject-wise single-cell simulations of neuronal response to a DC of various amplitudes — i.e., we estimated the f-I curves (Fig. 5A–B). The areas under curve (AUC) were 6.7% and 16.1% smaller in the SCZ compared to HC when data from PFC or ACC, respectively, was used (Fig. 5A–B). Since firing rate is inversely proportional to the membrane capacitance in the LIAF model (assuming a fixed leak conductance), we implemented these data as our models of SCZ as a change of membrane capacitance parameter: the membrane capacitances of all excitatory neurons (EO, ES, ED, EP, EP2, ESD, EDD populations) were 19.2% or 7.2% (PFC and ACC, respectively) larger in the SCZ case compared to HC.

**Figure 5.**
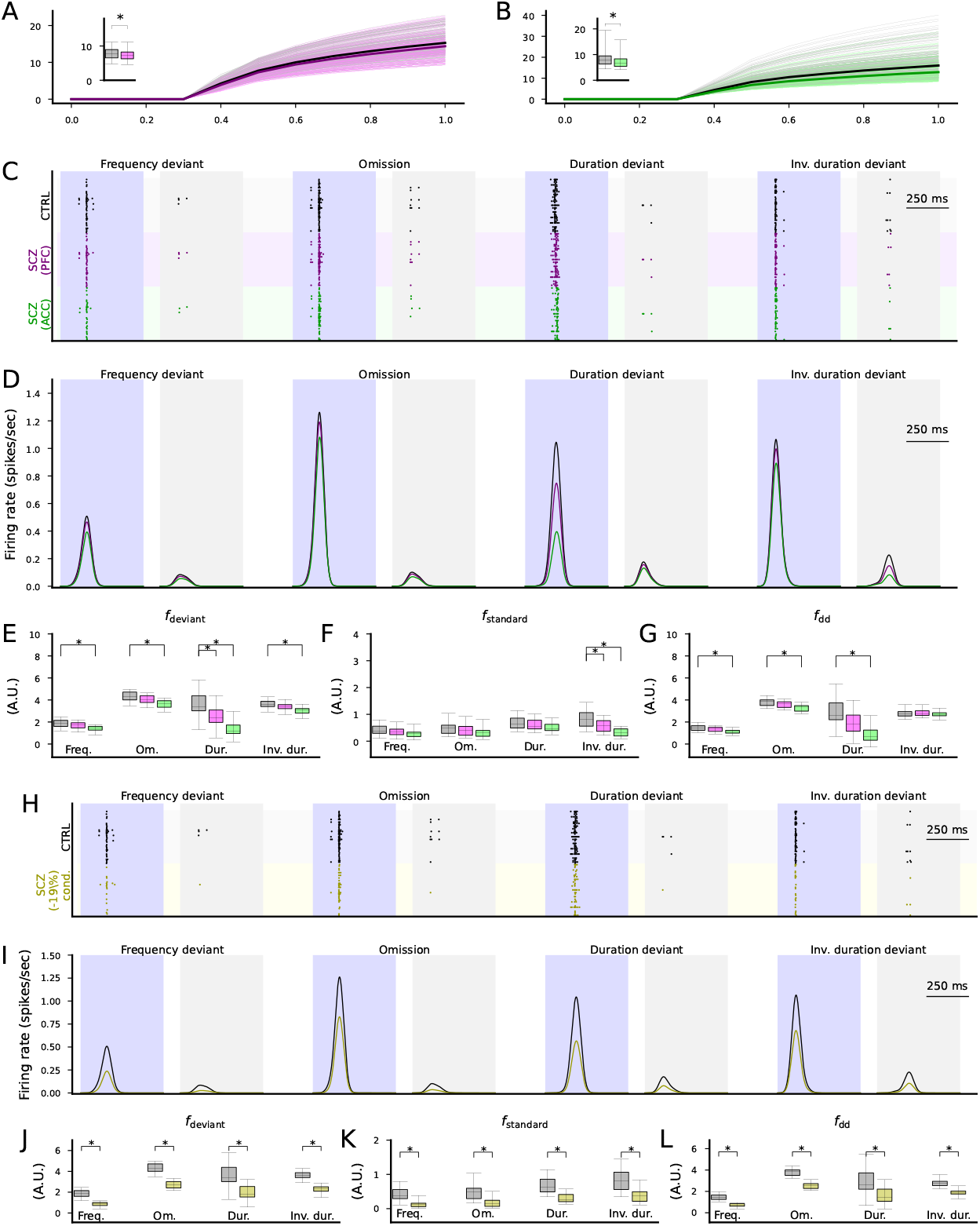
Altered expression of ion-channel-encoding genes and decreased spine density as measured postmortem in SCZ cortex can impair cortical novelty detection. **A–B**: The f-I curves of the layer V pyramidal cells where ion-channel conductances were adjusted based on post-mortem RNA expression data in the PFC (A) or ACC (B). Dim gray curves represent control subjects, and dim pink (A) and dim green (B) curves represent SCZ subjects. The thick curves represent the median f-I curves across the populations (black: control subjects, pink/green: SCZ subjects). Insets: box plot of the AUC across the control (gray) or SCZ (pink/green) subjects. **C**: Spike trains of the cortical output population in response to the four protocols (frequency deviant, omission of stimulus, duration deviant, inverse duration deviant) in the CTRL condition (black) and when the membrane capacitance was adapted to SCZ sample simulations according to ionchannel expression data from the PFC (purple) or ACC (green). **D**: The firing rate curves (smoothed with a Gaussian with an SD of 25 ms) of the spike train data of the three conditions, averaged across the 16 acceptable models. All simulations showed a trend where CTRL response *>* SCZ PFC response *>* SCZ ACC response. **E–F**: The distribution of the total firing rates in response to each deviant (E) or standard (F) stimulus (from 50 ms before the stimulus onset until 450 ms after the stimulus onset) in the four protocols across the 16 acceptable models. The asterisks show the statistically significant differences between the CTRL and SCZ (PFC or ACC; U-test, p*<*0.05/4). **G**: The distribution of the novelty detection index calculated from the data of (E) and (F). The asterisks show the statistically significant differences between the CTRL and SCZ (U-test, p*<*0.05/4). **H**: Spike trains of the cortical output population in response to the four protocols in the CTRL condition (black) and when the excitatory synaptic currents were decreased by 19% to emulate decreased spine density in SCZ (yellow), as reported in auditory cortex [Shelton et al., 2015]. **I–L**: The experiments of (D)–(G) repeated for CTRL vs SCZ (−19% synaptic conductance).

We then performed the network simulations with the four MMN protocols using the altered membrane capacitances (Fig. 5C). The simulations showed a decreased response of the output population to both standards and deviants (Fig. 5D). The responses to deviants of all types (Fig. 5E) and the responses to standards in the inverse duration deviant protocol (Fig. 5F) were significantly (p*<*0.05/4) decreased in amplitude for ACC, and the deviance detection index was significantly decreased in amplitude in the frequency (p=0.00059), omission (p=0.00029) and duration deviant (p=1.3·10^*−*6^) protocols but not the inverse duration deviant protocol (p=0.65) for ACC (Fig. 5G). For PFC, the response to deviants in the duration deviant protocol (Fig. 5E) and the response to standards in the inverse duration deviant protocol (Fig. 5F) were significantly (p*<*0.05/4) decreased in amplitude, but the deviance detection index was not significantly affected (Fig. 5G).

We also performed simulations of decreased synaptic density in cortical excitatory neurons (EO, ES, ED, EP, EP2, ESD, EDD populations). Reducing excitatory synaptic conductances by 19% decreased response amplitudes (Fig. 5H–K) and significantly reduced the deviance detection index (Fig. 5L) across all protocols (p=6.3·10^*−*8^ in the frequency deviant and omission, p=0.00044 in the duration deviant, and p=1.3·10^*−*7^ in the inverse duration deviant protocol; Fig. 5L).

### 3.5 Novelty detection as a subcortical phenomenon: Effects of cortical SCZ-associated cellular changes on the cortical projection of the novelty detection

Although animal studies suggest that auditory novelty detection is stronger in cortical compared to subcortical regions [Lao-Rodríguez et al., 2024], there is evidence of stimulus-specific adaptation (SSA) and/or auditory novelty detecting neurons in subcortical regions too, inferior colliculus and thalamus in particular [Anderson et al., 2009, Pérez-González et al., 2005, Reches and Gutfreund, 2008, Malmierca et al., 2009]. Here, we explore the consequences of SCZ-associated cellular-level phenotypes on the predicted novelty detection under the assumption that *all modelled neuron populations are subcortical neurons*. Namely, the intrinsic properties and the functionality of our novelty detecting network for auditory stimuli could match with a colliculo-thalamic feed-forward network where the output population corresponds to a population in the thalamic MGB while the other populations correspond to neuronal populations in the IC (see Discussion). However, the effects of SCZ diagnosis on cellular-level properties (such as expression of ion channels and spine density) as well as circuit abnormalities are reported more often in the cortex than subcortical regions (cf. [Lewis et al., 2009, Berdenis van Berlekom et al., 2020, Duncan et al., 2025]). Likewise, the SCZ-associated alterations we used in the experiments of Fig. 5 were observed in cortical rather than subcortical areas. These data thus only allow us to study how the cortical projection, and possibly post-processing, of the novelty detecting neurons are affected by SCZ although the predicted subcortical dynamics of the novelty detection might occur identically in SCZ compared to HC. Therefore, we here introduce an additional, cortex-like output population that allows us to make model predictions based on these data.

We connected the previously described output population (EO) to a cortical output population (CO) of the same size (N=40). We used membrane properties that matched the multicompartmental Hay model of layer V pyramidal cells [Hay et al., 2011], namely, a membrane capacitance of 580 pF and a leak conductance of 4 nS (Fig. 6A) — both parameters exhibited a 30% standard deviation within the neuron population centered at these values, similar to other populations (see Section 3.1). To mimic in vivo variability, random membrane currents induced spontaneous fluctuations and spiking. We adjusted the connection strengths from the EO to the CO population and the amplitude of the random membrane currents to match with experimental data on the AP frequency during a response to standards (∼ 8 spikes/sec) and oddballs (∼ 20 spikes/sec) [Ulanovsky et al., 2003]) — the best fit to data was obtained by *g*_*AMP A*_ = 3 µS and *I*_*noise,SD*_ = 1.75 nA (Fig. 6B). We ran the resulting network model and assessed the activity of the cortical output population (Fig. 6C) in the different stimulus protocols with all 16 acceptable parameter sets (see Section 3.1). The model provided a strong signal for the novel stimuli in all four stimulus protocols compared to the standards (Fig. 6D). In the following, we use the cortical output as the final output affecting the EEG signal and test the effects of different SCZ-associated cellular-level alterations on this output.

**Figure 6.**
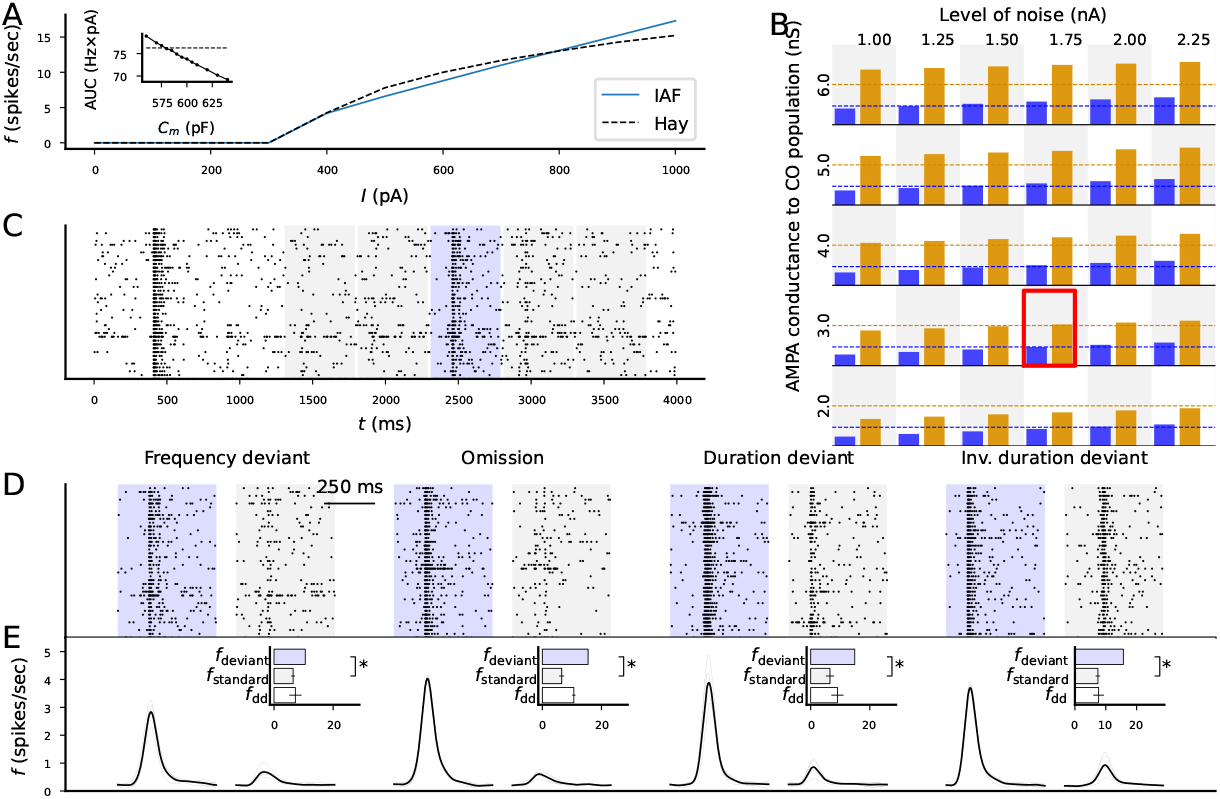
Introduction of a cortical output population and fitting its properties to electrophysiological data from animal oddball experiments. **A**: Best-fit IAF model to the f-I curve of the Hay model [Hay et al., 2011] of cortical pyramidal neurons. Inset: AUC (y-axis) of IAF models with different membrane capacitances (x-axis). The best fit was obtained by *τ* =580 ms. **B**: Firing rates of the cortical output population in response to standards (blue) and deviants (orange) when different levels of noise and connection strengths from the (subcortical) output population were used. Dashed lines indicate the data recorded from cats [Ulanovsky et al., 2003]. Best fit was obtained by a noise level of 1.75 nA and a connection strength of 3 µS. **C**: Example spike train of the cortical output population during the frequency deviant protocol. The blue shaded area represents the time interval from which the response to deviants were determined whereas the gray areas represent the time intervals from which the response to standards were determined. The firing rate from the four responses to standards were averaged. **D**: Example spike trains of the cortical output population during the deviant and the first standard after the deviant in all four protocols. **E**: Firing rate curves averaged across the 16 models and 20 seeds in response to deviants (left) and standards (right) in all four protocols. The top and middle bars show the mean and SD of the firing rates across 20 random number seeds in response to deviants (blue) and standards (gray). The bottom bar shows the mean and SD of the deviance detection index (white).

We next fit the membrane capacitance in the IAF model to each of the subject-wise adapted simulations of the Hay model (Fig. 7A–B) in a similar way as done for the default Hay model in the experiment of Fig. 6A. These fits suggested that the membrane capactitance is on average 631 or 678 pF in SCZ according to the PFC or ACC data, respectively (Fig. 7A–B insets), compared to the 580 pF fitted for the default Hay model. The IAF model simulations based on these data are illustrated in Fig. 7C–D.

**Figure 7.**
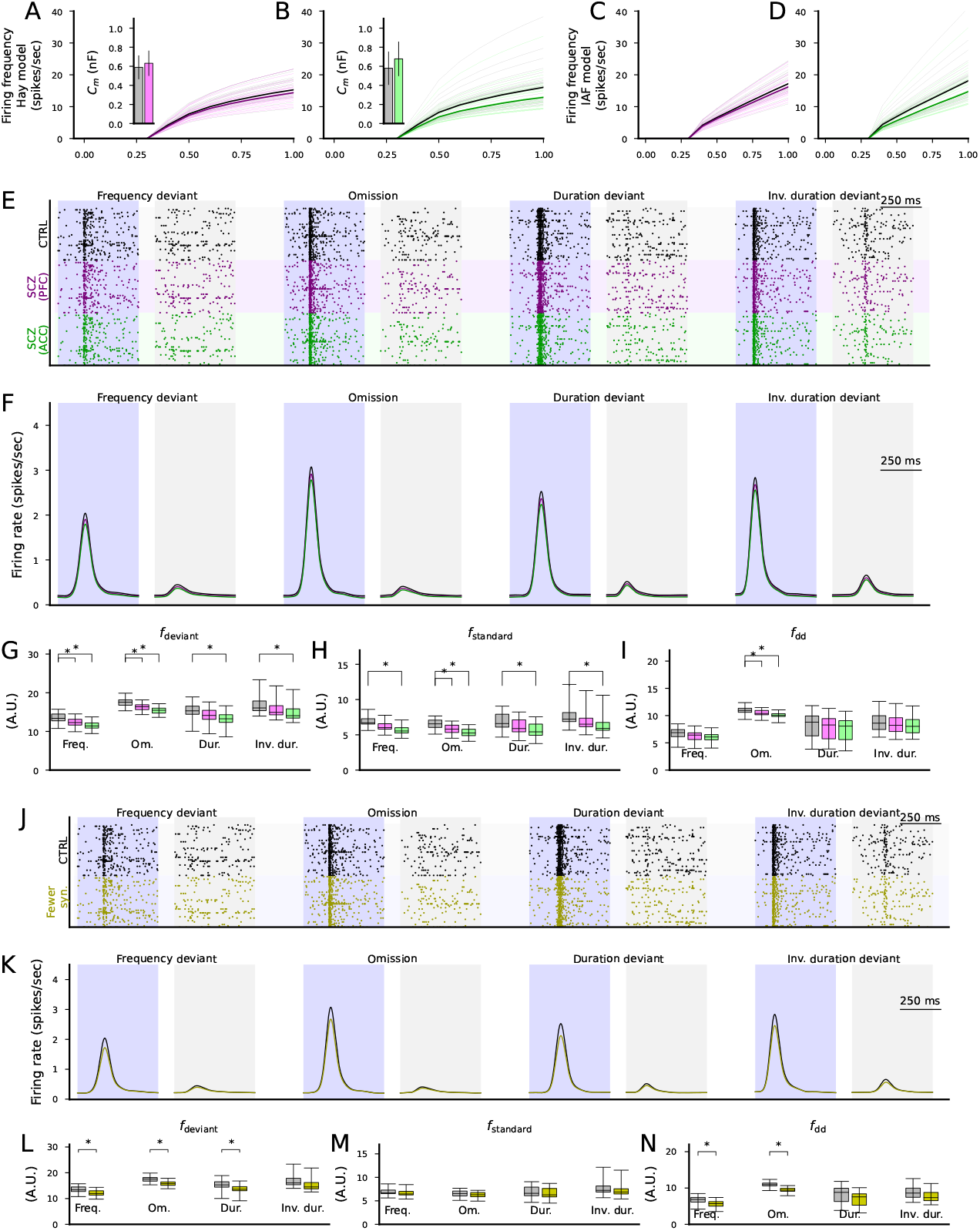
Altered expression of ion-channel-encoding genes and decreased spine density as measured postmortem in SCZ cortex can impair cortical novelty detection. **A–B**: The f-I curves of the layer V pyramidal cells where ion-channel conductances were adjusted based on post-mortem RNA expression data in the PFC (A) or ACC (B). Dim gray curves represent control subjects, and dim pink (A) and dim green (B) curves represent SCZ subjects. The thick curves represent the median f-I curves across the populations (black: control subjects, pink/green: SCZ subjects). Insets: box plot of the AUC across the control (gray) or SCZ (pink/green) subjects. **C–D**: The f-I curves of best-fit IAF models fitted to the data from (A)–(B). **E**: Spike trains of the cortical output population in response to the four protocols (frequency deviant, omission of stimulus, duration deviant, inverse duration deviant) in the CTRL condition (black) and when the membrane capacitance was adapted to SCZ sample simulations according to ion-channel expression data from the PFC (purple) or ACC (green). **F**: The firing rate curves (smoothed with a Gaussian with an SD of 25 ms) of the spike train data of the three conditions, averaged across the 16 acceptable models. All simulations showed a trend where CTRL response *>* SCZ PFC response *>* SCZ ACC response. **G–H**: The distribution of the total firing rates in response to each deviant (G) or standard stimulus (from 50 ms before the stimulus onset until 450 ms after the stimulus onset) in the four protocols across the 16 acceptable models. The asterisks show the statistically significant differences between the CTRL and SCZ (PFC or ACC; U-test, p*<*0.05/4). **I**: The distribution of the novelty detection index calculated from the data of (G) and (H). The asterisks show the statistically significant differences between the CTRL and SCZ (U-test, p*<*0.05/4). **J**: Spike trains of the cortical output population in response to the four protocols in the CTRL condition (black) and when the excitatory synaptic currents were decreased by 19% to emulate decreased spine density in SCZ (yellow). **K–N**: The experiments of (F)–(I) repeated for CTRL vs SCZ (−19% synaptic conductance).

We next simulated the novelty detecting network with the cortical output population whose membrane capacitance was set to 580 pF (CTRL), 631 pF (SCZ, PFC) or 678 pF (SCZ, ACC). The simulations showed a decreased response of the cortical output population to both standards and deviants (Fig. 7E–F). For ACC, responses to both deviants and standards were significantly weaker in SCZ in all protocols, whereas for PFC, the responses to deviants were significantly weaker in the frequency and stimulus omission protocols, and the reponses to standards were significantly weaker in the stimulus omission protocol (Fig. 7G–H). In both brain areas, the deviance detection index was significantly decreased in the stimulus omission protocol (p=0.00019 in ACC and p=0.012 in PFC) but not other protocols (Fig. 7I).

Finally, we simulated the effect of decreased synaptic conductance of the putative thalamocortical synapses on the novelty detection. Similar to the experiments where the synaptic conductances of all excitatory populations were decreased (Fig. 5E–F), the decrease in the synaptic conductances of the cortical output population only decreased the response to both standards and deviants (Fig. 7J–K). However, the simulations suggested that the decrease in the response to deviants was statistically significant in the frequency deviant, stimulus omission and the duration deviant protocols (Fig. 7L) while the decrease in the response to standards was not statistically significant in any protocol (Fig. 7M). The deviance detection index was significantly decreased in the frequency deviant and the omission protocol (p=0.0038 and p=2.2×10^*−*6^, respectively) but not in the duration deviant or inverse duration deviant protocols (Fig. 7N).

Taken together, our simulations suggest that even if the deviance detection takes place in the subcortical areas, SCZ-like alterations in the excitatory cortical population receiving the subcortical signals can decrease the novelty detection amplitude. SCZ-associated ion-channel expression differences as measured in the PFC did not significantly affect the deviance detection index, while those measured in the ACC significantly decreased the deviance detection index in the frequency deviant protocol, and the decreased synaptic conductance (to simulate the decrease in numbers of post-synaptic spines in SCZ) decreased the deviance detection index in the frequency deviant and omission protocols.

## 4 Discussion

We developed a spiking network that can detect four types of deviations from standard stimuli: frequency deviants (activating different pitch-sensitive neurons than standards), stimulus omissions, duration deviants (longer than standards), and inverse duration deviants (shorter than standards). We showed that the network performs robustly in a series of experiments where the stimuli or the neuron properties were mildly altered. We also explored the behaviour of the network in the presence of the SCZ-associated cellular-level alterations, namely, an altered expression of ion-channel-encoding genes and an altered density of postsynaptic spines. Our analysis suggests that both a 19% decrease in EPSCs to the excitatory neurons as well as changes in ion-channel-encoding gene expression that decreases the neuronal excitability can impair the novelty detection, but that the former had stronger effects than the latter. This observation was made both when interpreting our novelty detection network to operate cortically (where all excitatory neurons were assumed to be affected by the SCZ-associated cellular-level alterations) or mainly subcortically (where only the cortical output population was affected), although the former interpretation led to larger effects altogether.

A key aspect of this study lies in its assumptions and modelling choices. Our model was built in a simplified, yet biophysically meaningful manner, and the modelling choices were primarily made to optimize the model for auditory novel detections. The suggested neuron populations may therefore represent neurons of multiple neuron types (especially the inhibitory populations) and even neurons at different brain areas rather than a specific anatomical region. However, our modelling choices are not, as far as we are aware, inconsistent with the existing literature. Here, we discuss each of our assumptions and modelling choices in the light of the previous findings and models as well as the alternatives.

1. **Short-term depressing excitatory synapses between neurons in the auditory pathway**. A central assumption in our model is that novelty detection relies on stimulus-specific adaptation (SSA). Apart from the very upstream cochlear nucleus [Ayala et al., 2013], SSA has been observed in most brain areas in the auditory pathway, including inferior colliculus (IC) [Ayala et al., 2013], superior olivary complex (SOC) [Finlayson and Adam, 1997], thalamus [Anderson et al., 2009], and the auditory cortex [Ulanovsky et al., 2003]. It is thus natural to hypothesize that neurons encoding for auditory novelty receive direct or indirect inputs from neurons exhibiting SSA. The mechanisms underlying SSA have not been confirmed, but presynaptic short-term depression has been suggested as a central mechanism for adaptation occurring at time scales larger than 100 ms [Wehr and Zador, 2005]. We thus built our network to rely on models of presynaptic short-term depression [Tsodyks and Markram, 1997, Wang, 1999].
2. **Duration-sensitive neurons**. In this work, we assumed that the frequency-tuned neurons are divided into two subgroups: those that only respond to long stimuli and those that respond to both. Such long-pass firing behaviours have been observed in vivo [Ehrlich et al., 1997, Brand et al., 2000, Sayegh et al., 2011], but there is evidence also of neurons that exclusively respond to short, medium-duration, or long stimuli [Ehrlich et al., 1997, Aubie et al., 2014]. A previous theoretical study suggested that varying the membrane time constant can explain the lag in response to auditory stimulus [Aubie et al., 2012] — we thus implemented the long-pass duration sensitivity by allowing different membrane time constants (capacitances) to the late-responding populations (ESD, EDD) that were fitted to optimize the deviance detection. However, if the exclusively duration-sensitive neurons (as observed in, e.g., [Ehrlich et al., 1997, Aubie et al., 2014]) are dominant in the auditory pathway, the detection of duration deviants could be implemented in a simpler way where each neuron population responds not only in a frequency-selective manner but in a frequency and duration-selective manner. This would mean that the detection of duration deviants could be conceptually identical to the detection of frequency deviants in our modelling framework: repetitive introduction of tones with a particular frequency and duration depresses the synapses from neurons coding this frequency and duration, and a change in either frequency or duration of the tone will activate other neurons whose out-going synapses are not short-term depressed.
3. **Existence of neurons that fire phase-locked to a rate with which standards are presented**. To permit detection of omitted standards, the brain has to know what to predict and, importantly, when to predict. Previous models (e.g., [Wacongne et al., 2012, Lieder et al., 2013]) have circumvented this problem by assuming a kind of “cache”, that is, a neuron population that saves the memory of the recently income stimulus and replays it with a delay. While there is no direct evidence of such behaviour or suggested mechanisms how this memory-trace firing could emerge, we are not aware of direct evidence for our assumption of delta-frequency phase-locked neurons or inputs from such neurons in the auditory pathway either. However, there are findings that indirectly support our assumption. Namely, there are cortical and subcortical neurons that exhibit robust pacemaking activity (reviewed in [Bean, 2024]). For example, the subthalamic neurons showed spontaneous rhythmic firing in the delta-theta frequency band in vitro [Bevan and Wilson, 1999]. Likewise, thalamic relay neurons are well known for their spontaneous rhythmic generation of bursts of action potentials in a delta-band frequency [Timofeev et al., 2012]. It has also been observed that low-frequency cortical oscillations can become phase-locked by introduction of a rhythmic stimulus repeated at a 1.5 Hz rate [Ten Oever et al., 2017]. It is thus natural to hypothesize that standards repeated at a nearby stimulus presentation rate, 2.0 Hz, also make certain population fire at this rate. To argue for the plausibility of our network architecture, we here proposed that the phase-locking to a rhythmic auditory stimulus may emerge in the auditory pathway as a consequence of STDP and phase-locked synaptic inputs from elsewhere in the brain (Fig. 4). However, it remains to be shown that there are significant neuron populations phase-locked to these oscillations at sufficiently many phase intervals.
4. **The role of the output population**. The introduction of the output population in all our simulations serves the purpose of showing that it is possible for a single population in the brain to encode auditory novelty of many types in a centralized manner. However, this is not a necessity for the induction of the MMN signal or for shifting the animal’s attention toward the cause of the auditory novelty, nor is there evidence that the brain operates this way. The assumption of the existence of such a centralized population is, however, useful for quantifying the network’s capacity for novelty detection.

In our SCZ-applied simulations, we assumed that all neurons were either cortical or subcortical, but, as discussed above, they could be distributed across different cortical and subcortical areas as well. However, one of the brain areas that would match with our modelled novelty-detecting network both by anatomy and function is the IC. IC receives direct inputs from the upstream auditory areas, i.e. the cochlear nucleus and the superior olivary complex [Adams, 1979, Grothe et al., 1994]. Approximately 30% of IC neurons are inhibitory, and they connect to both local IC neurons and neurons in other brain areas, including the thalamus [Peruzzi et al., 1997, Oberle et al., 2023]. Importantly, IC exhibits strong, rapid inhibition in the form of binaural interaction that is important for sound localization [Sanes et al., 1998] — these same network mechanisms could underlie the inhibition of the phase-locked population needed for omission detection in our network. IC neurons also display a wide (more than 10-fold) diversity of excitability-related properties such as time to spike in response to EPSP onset and threshold stimulus intensity [Tan and Borst, 2007], which could give rise to populations responding to long but not short auditory stimulus such as the ES and ED populations described in our model. The phase locking to the 2-Hz stimulus could possibly be mediated by cortico-collicular pathway [Lesicko and Geffen, 2022], but further research is needed to test this. On the other hand, the requirement for phase-locking in the 2-Hz frequency and the possible requirement of STDP to attain it may fit better to cortical than subcortical circuits.

The existence of neurons in the auditory pathway receiving inputs that are phase-locked to different phases of the stimulus rate is an assumption that is needed in our framework for the detection of omitted stimuli. In this work, we showed that the timing of these neurons does not need to be perfectly aligned for a successful omission detection (Fig. S6D–E), and that the network also tolerates small and medium synaptic delays (Fig. 3F–G). We also showed that the phase locking can arise in response to the rhythmic auditory stimulus given synaptic inputs obeying an STDP rule from other neuron populations that are phase locked (Fig. 4) — if this is the case, the deviance detection of omitted stimuli should be weak in the beginning of the MMN experiment, since STDP typically takes many minutes to become fully effective (cf. [Bi and Poo, 1998]). Although the experiment of Fig. 4 still required a population to be phase locked to the possible stimulus rate, it showed that this requirement applied only to the inputs of these neurons in the auditory pathway, and thus fewer neurons in the auditory pathway itself would be needed to attain this form of deviance detection. The requirement of many phases of the phase locking could also be avoided by an architecture permitting synfire chains — these architectures may also be formed through plasticity rules [Zheng and Triesch, 2014]. The phase-locking experiment of (Fig. 4) has also been previously performed using mean field modelling [Luz and Shamir, 2016], albeit in different frequency ranges than ours. Note that the learning of rhythmic inputs may also be attained through alternative ways, such as those employed in [Socolovsky and Shamir, 2021]. All these theoretical analyses point to the eligibility of the omission detection mechanisms we propose; however, experimental research is needed to confirm the existence of phase-locked inputs to the auditory pathway neurons.

Most of our simulation experiments were carried out using the four MMN protocols that were used in the fitting of the model parameters: short standard to short deviant tone, omission after short standard tones, short standard to long standard tone, and long standard to short standard tone. We also tested that the network successfully detected novelty in all other combinations of the four types of tones that were introduced in a roving paradigm-like manner and omissions from the other standard sequences (Fig. 3A–C). We can expect that the model would function equally well if additional tones were added as alternative stimulus frequencies in addition to the currently implemented “standard” and “deviant” tones. However, the network would not detect more intricate patterns such as the cascade or many-standards control sequences [Harms et al., 2014] without a more thorough redesign of the network.

SCZ is associated with many cellular-level phenotypes pertaining to neuron morphology and electrophysiology. In this work, we considered two central alterations affecting pyramidal neuron behaviour in the cortex, namely, a 19% loss of synaptic spines [Shelton et al., 2015] and an altered expression of ion-channel-encoding genes [Hoffman et al., 2019]. Beyond these, NMDA receptor hypofunction has long been proposed as a central mechanism underlying MMN deficits, supported by human pharmacological studies where NMDA antagonists such as ketamine reliably reduce MMN amplitudes [Umbricht et al., 2000, Javitt et al., 2008]. Dysfunction of GABAergic interneurons, particularly parvalbumin-positive cells, has also been associated with disrupted excitatory-inhibitory balance and abnormal auditory oscillations [Gonzalez-Burgos and Lewis, 2012, Metzner et al., 2019]. These alterations could interact with the excitatory deficits modelled here, potentially compounding their effects. Other reported phenotypes include altered dendritic morphology as well as oligodendrocyte- and myelin-related abnormalities; these changes could further degrade the temporal precision and conduction necessary for omission and duration-deviant responses [Kalus et al., 2000, Takahashi et al., 2011, Valdés-Tovar et al., 2022].

At the systems level, empirical findings suggest that not all types of auditory deviants are equally affected in SCZ. Several studies report stronger MMN impairments for duration deviants compared to frequency deviants [Todd et al., 2012, Erickson et al., 2016], a dissociation that resonates with our model’s prediction that different deviant types may have different vulnerabilities to cellular-level alterations. Moreover, the ongoing debate about the relative contributions of cortical versus subcortical structures to MMN generation [Kraus et al., 1992, Grimm and Escera, 2012] is directly reflected in our dual interpretations of the network, which yielded converging but distinct predictions about SCZ-associated impairments.

Our computational analysis is valuable in providing mechanistic in silico predictions against which future observations can be reflected. Pharmacological manipulations that differentially perturb excitatory drive and NMDA receptor function could clarify whether MMN deficits are more strongly driven by excitability loss, synaptic adaptation, or receptor hypofunction. Further validation of our model could be obtained through in vitro slice recordings that test how spine loss and excitability changes shape SSA and omission responses at the circuit level. Finally, patient stratification studies — comparing SCZ subgroups or genetic risk carriers — may reveal whether particular forms of MMN deficits (e.g., omissions versus duration deviants) track with specific cellular phenotypes. Since our model can be tuned to individual gene expression data, it enables personalized predictions of auditory novelty detection deficits and may help disentangle the heterogeneity of SCZ. Given experimental support for the central mechanisms and an identification of the neuron classes behind the proposed populations, our model could then be reused with targeted genetic and structural psychiatric data to better explain the mechanisms of MMN deficits in SCZ, paving way for an improved understanding of the pathophysiology of the disorder.

## 5 Acknowledgements

## 6 Author contributions

Conceptualization: AE, CM, TE, MLL, GTE, OAA, TMM. Data curation: ABP, IA, SD, TE, AS. Investigation: AE, JFK, TMM. Project administration: TMM. Resources: SD, TE, OAA. Writing original draft: AE, TMM. Writing, review and editing: AE, JFK, CM, ABP, IA, SD, TE, AS, MLL, GTE, OAA, TMM.

## Supplementary figures and tables

**Figure S1.**
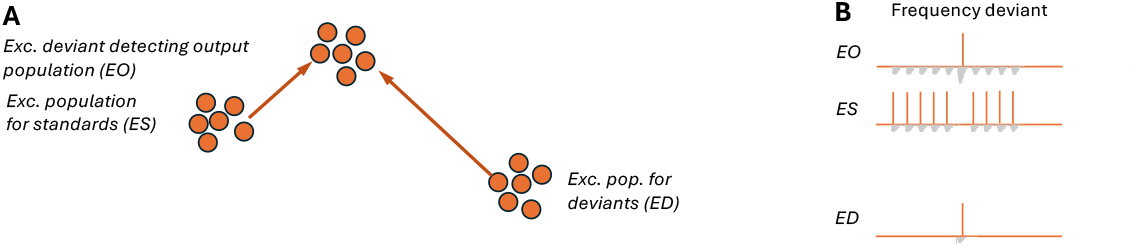
Illustration of a the hypothesized mechanisms of deviance detection for the frequency deviants and the part of the network required for this. **A**: Illustration of the part of the network. ES and ED populations project to the EO population with excitatory, short-term depressing synaptic connections. **B**: Illustration of the expected response of the network to a sequence of standard tones, where one of the stimuli is replaced by a deviant tone. The gray EPSCs illustrate the EPSCs received by the neuronal population. The EO population requires **a large EPSC, caused by a first (but not successive) stimulus of the corresponding tone**, to fire.

**Figure S2.**
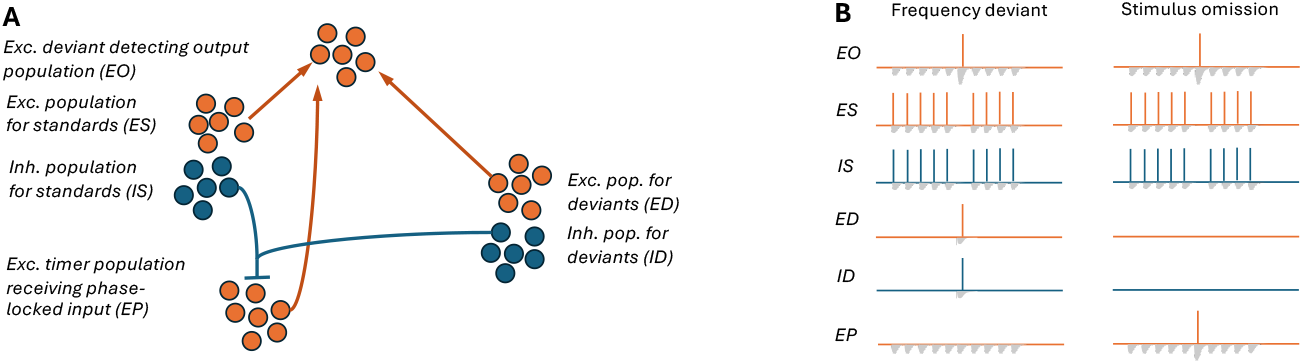
Illustration of a the hypothesized mechanisms of deviance detection for the frequency deviants and the omitted stimuli and the part of the network required for this. **A**: Illustration of the part of the network. In addition to the network in Fig. S1, IS and ID populations project to the EP population with inhibitory synaptic connections. The EP population projects to the EO population with excitatory, short-term depressing synaptic connections. **B**: The first column shows the responses to the sequence discussed in Fig. S1, while the second column illustrates the expected response of the network to a sequence of standard tones where one of the stimuli is omitted. The EO population requires either a large EPSC caused by a first stimulus of the corresponding tone **or a large EPSC from the EP population** to fire. The EP is driven by rhythmic inputs arriving at a certain phase of the stimulus rate (2 Hz in this work) but is typically inhibited by auditory stimuli arriving at the same time.

**Table S1:**
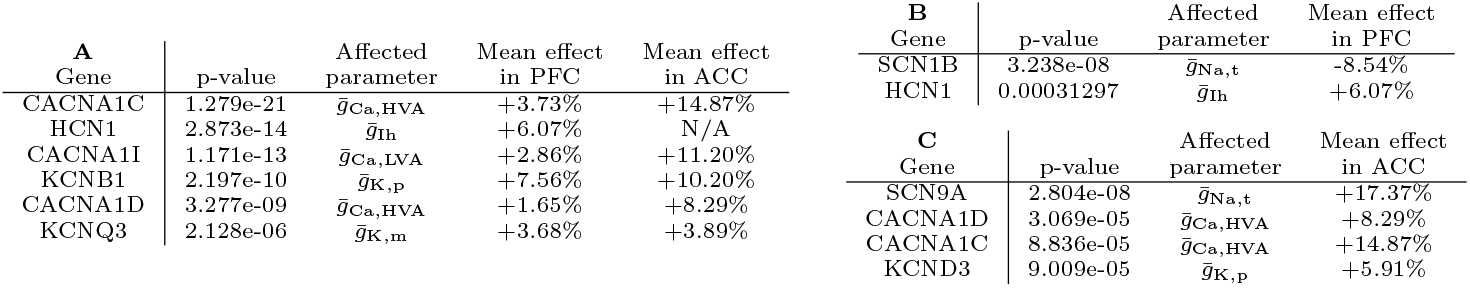
Table of ion channel-encoding genes associated with SCZ according to GWAS [Trubetskoy et al., 2022] or postmortem expression data from the PFC (B) or ACC (C).

**Figure S3.**
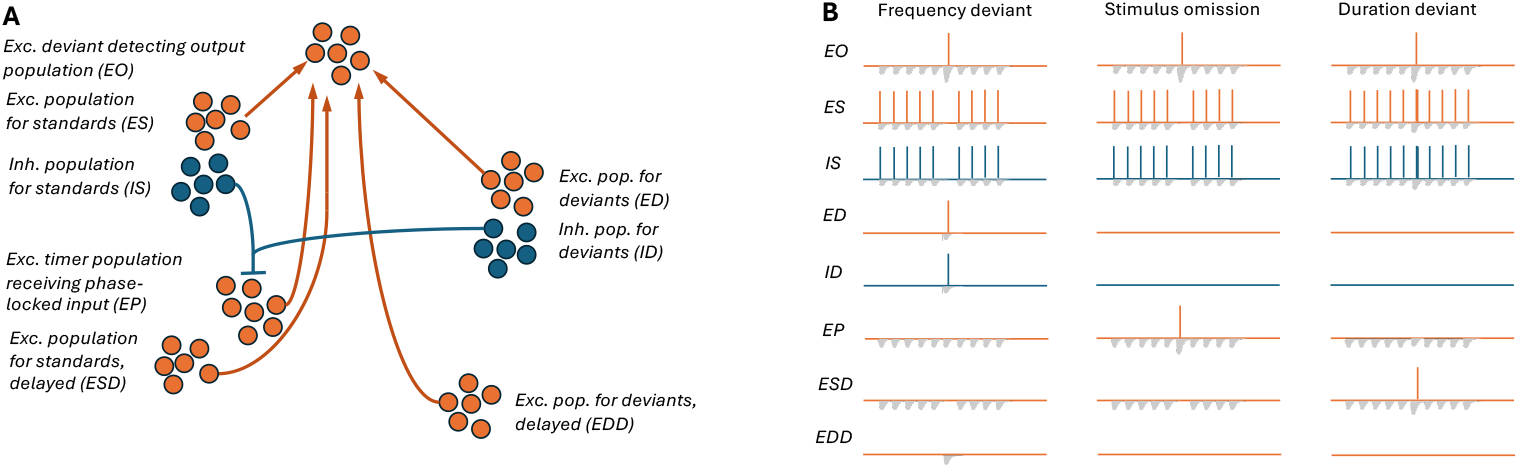
Illustration of a the hypothesized mechanisms of deviance detection for the frequency and duration deviants and the omitted stimuli and the part of the network required for this. **A**: Illustration of the part of the network. In addition to the network in Fig. S2, ESD and EDD populations project to the EO population with excitatory, short-term depressing synaptic connections. **B**: The first two columns show the responses to the sequences discussed in Fig. S2, while the third column illustrates the expected response of the network to a sequence of standard tones where one of the stimuli is replaced by a longer stimulus. The EO population requires either a large EPSC caused by a first stimulus of the corresponding tone, a large EPSC from the EP population, **or an EPSC from both ES/ED and ESD/EDD** to fire. The ESD and EDD populations are activated by the same auditory stimuli as ES and ED, but they require a longer presentation of the stimulus (i.e., a longer integration of the corresponding inputs) to fire.

**Figure S4.**
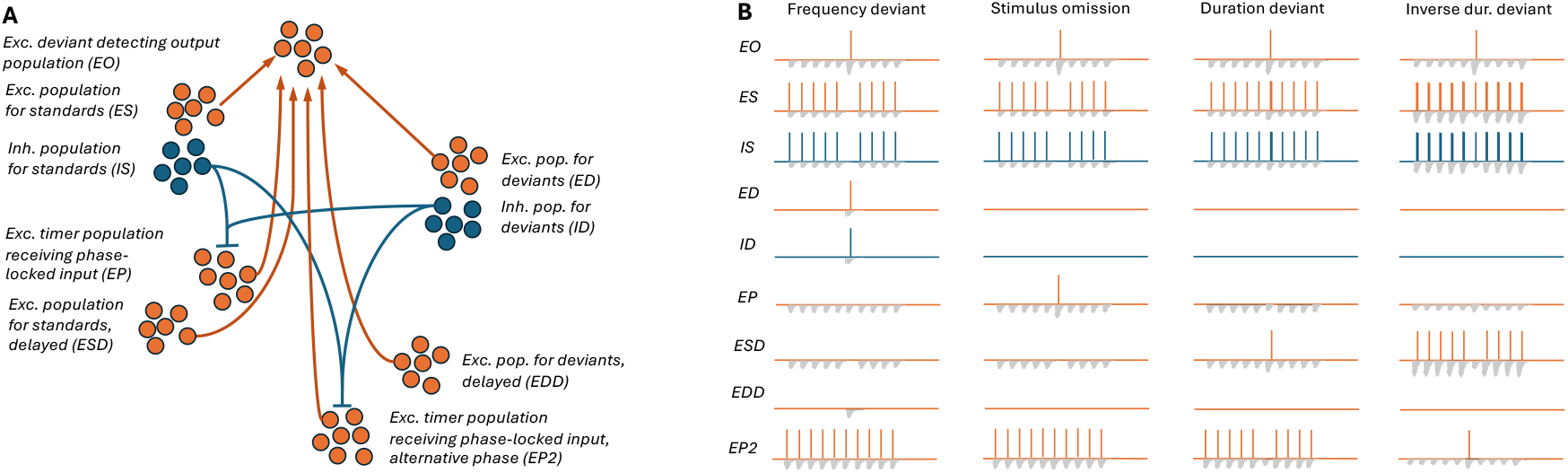
Illustration of a the hypothesized mechanisms of deviance detection for all deviants. **A**: Illustration of the part of the network. In addition to the network in Fig. S3, IS and ID populations project to the EP2 population with inhibitory synaptic connections. Similar to the EP population, the EP2 population projects to the EO population with excitatory, short-term depressing synaptic connections. **B**: The first three columns show the responses to the sequences discussed in Fig. S3, while the fourth column illustrates the expected response of the network to a sequence of long standard tones where one of the stimuli is replaced by a shorter stimulus. The EO population requires either a large EPSC caused by a first stimulus of the corresponding tone, a large EPSC from the EP **or EP2** population, an EPSC from both ES/ED and ESD/EDD to fire. The EP2 is driven by rhythmic inputs arriving at a certain phase (prior to the preferred phase of the EP population) of the stimulus rate but is typically inhibited by long, auditory stimuli arriving at the same time. The short stimulus deviant allows EP2 to fire, causing EO to fire as well. The EP2 population fires typically in the other protocols (first to third column), but the inputs caused by these activations become short-term depressed and thus cause no firing of EO population.

**Figure S5.**
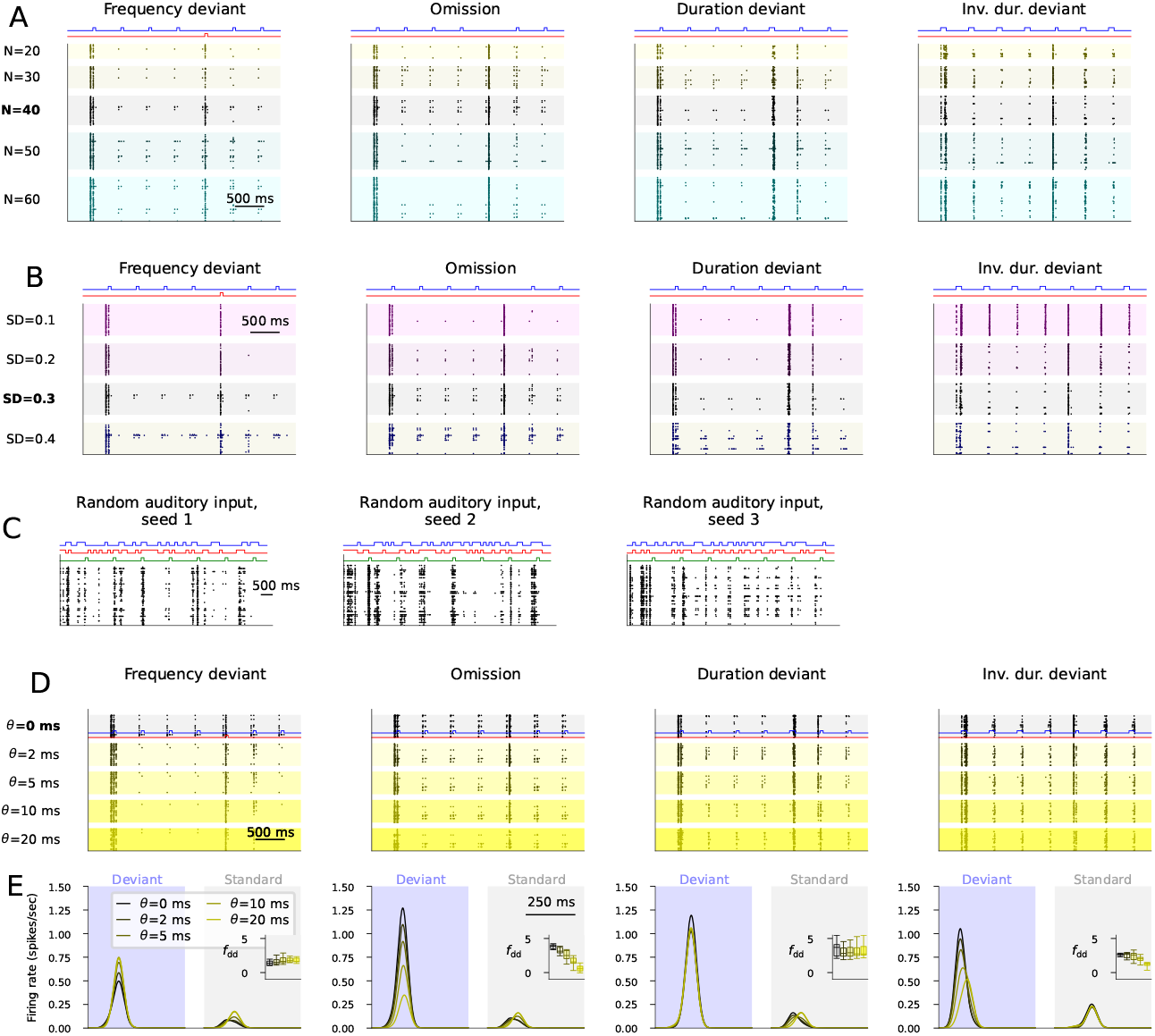
The deviance detection network is robust against changes in network size and intrinsic variability as well as against small jitter in the phase of the phase-locked populations. **A**: The population spike trains of the output population when different population sizes were used. The default parameter (N=40 neurons per population) is shaded in gray. In each experiment with an altered network size, all synaptic conductances were compensated by an inverse factor (i.e., when network size was changed from 40 to 60, synaptic conductances were divided by 1.5 from their default value). **B**: The population spike trains of the output population when different intrinsic variabilities of the membrane capacitance (*C*_*m*_) were used. The default parameter (SD(*τ*)=0.3 × mean(*τ*)) is shaded in gray. Slightly altered variabilities (SD=0.2, 0.4) resulted in successful deviance detection, but a network where all neurons within a population had exactly the same membrane capacitance (SD=0) resulted in a compromised detection of inverse duration deviants. **C**: An experiment where the auditory inputs are random does not show artifacts caused by the phase-locked populations. The three panels show the results from experiments performed with three different random number seeds. **D**: Experiments where the phase-locked neurons had a jitter of up to 0 (default), 2, 5, 10, or 20 ms around their expected time of firing. **E**: The experiments of (D) repeated for all 16 models and 10 repetitions with different random number seeds. The insets show the deviance detection indices for the jitter of 0 (left), 2, 5, 10, and 20 ms (right). The detection of omissions is compromised for medium (10 ms) and large (20 ms) jitter, and the detection of inverse duration deviants is impaired for large (20 ms) jitter, while the detection of frequency and duration deviants is unaffected by the jitter.

**Table S2:** Parameters and their ranges used in the grid search (A) and the acceptable parameter sets obtained (B). The default model parameters are printed in bold.

|  |  |  |  |  |  |  |  |
| --- | --- | --- | --- | --- | --- | --- | --- |
| P1: Stimulus amplitude (ES, IS, ED, ID, ESD, EDD) | 120 | 125 | 130 | 140 | 150 | (pA) |  |
| P2: Synaptic conductance from ES and ED to EO | 12.5 | 15.0 | 17.5 | 20.0 | 22.5 | (nS) |  |
| P3: Synaptic conductance from EP and EP2 to EO | 25.0 | 30.0 | 35.0 | 40.0 | 45.0 | (nS) |  |
| P4: Synaptic conductance from ESD and EDD to EO | 80.0 | 90.0 | 100.0 | 110.0 | 120.0 | 130.0 | 140.0 (nS) |
| P5: NMDA to AMPA ratio | 0.5 | 0.333 |  |  |  |  |  |
| P6: Synaptic conductance from IS and ID to EP and EP2 | 10.0 | 15.0 | 20.0 | 25.0 | 30.0 | 35.0 | (nS) |
| P7: Strength of depression $p_v$ | 0.9 | 0.95 | | | | | |
| P8: Average membrane capacitance of ESD and EDD | 200 | 250 | (pF) |  |  |  |  |

|  | P1 | P2 | P3 | P4 | P5 | P6 | P7 | P8 |
| --- | --- | --- | --- | --- | --- | --- | --- | --- |
| **Parameter set 1** | **150** | **17.5** | **30.0** | **80.0** | **0.333** | **35.0** | **0.95** | **250.0** |
| Parameter set 2 | 150 | 17.5 | 30.0 | 80.0 | 0.333 | 35.0 | 0.9 | 250.0 |
| Parameter set 3 | 150 | 17.5 | 30.0 | 80.0 | 0.333 | 30.0 | 0.95 | 250.0 |
| Parameter set 4 | 150 | 17.5 | 30.0 | 80.0 | 0.333 | 30.0 | 0.9 | 250.0 |
| Parameter set 5 | 150 | 17.5 | 30.0 | 80.0 | 0.333 | 25.0 | 0.95 | 250.0 |
| Parameter set 6 | 150 | 17.5 | 30.0 | 80.0 | 0.333 | 25.0 | 0.9 | 250.0 |
| Parameter set 7 | 150 | 17.5 | 30.0 | 80.0 | 0.333 | 20.0 | 0.9 | 250.0 |
| Parameter set 8 | 150 | 17.5 | 30.0 | 80.0 | 0.333 | 15.0 | 0.95 | 250.0 |
| Parameter set 9 | 150 | 17.5 | 30.0 | 80.0 | 0.333 | 15.0 | 0.9 | 250.0 |
| Parameter set 10 | 140 | 17.5 | 30.0 | 80.0 | 0.333 | 35.0 | 0.9 | 250.0 |
| Parameter set 11 | 140 | 17.5 | 30.0 | 80.0 | 0.333 | 30.0 | 0.9 | 250.0 |
| Parameter set 12 | 140 | 17.5 | 30.0 | 80.0 | 0.333 | 10.0 | 0.9 | 250.0 |
| Parameter set 13 | 125 | 17.5 | 40.0 | 140.0 | 0.333 | 25.0 | 0.9 | 250.0 |
| Parameter set 14 | 125 | 17.5 | 40.0 | 140.0 | 0.333 | 10.0 | 0.9 | 250.0 |
| Parameter set 15 | 125 | 17.5 | 40.0 | 100.0 | 0.333 | 25.0 | 0.9 | 200.0 |
| Parameter set 16 | 125 | 17.5 | 40.0 | 100.0 | 0.333 | 10.0 | 0.9 | 200.0 |

**Figure S6.**
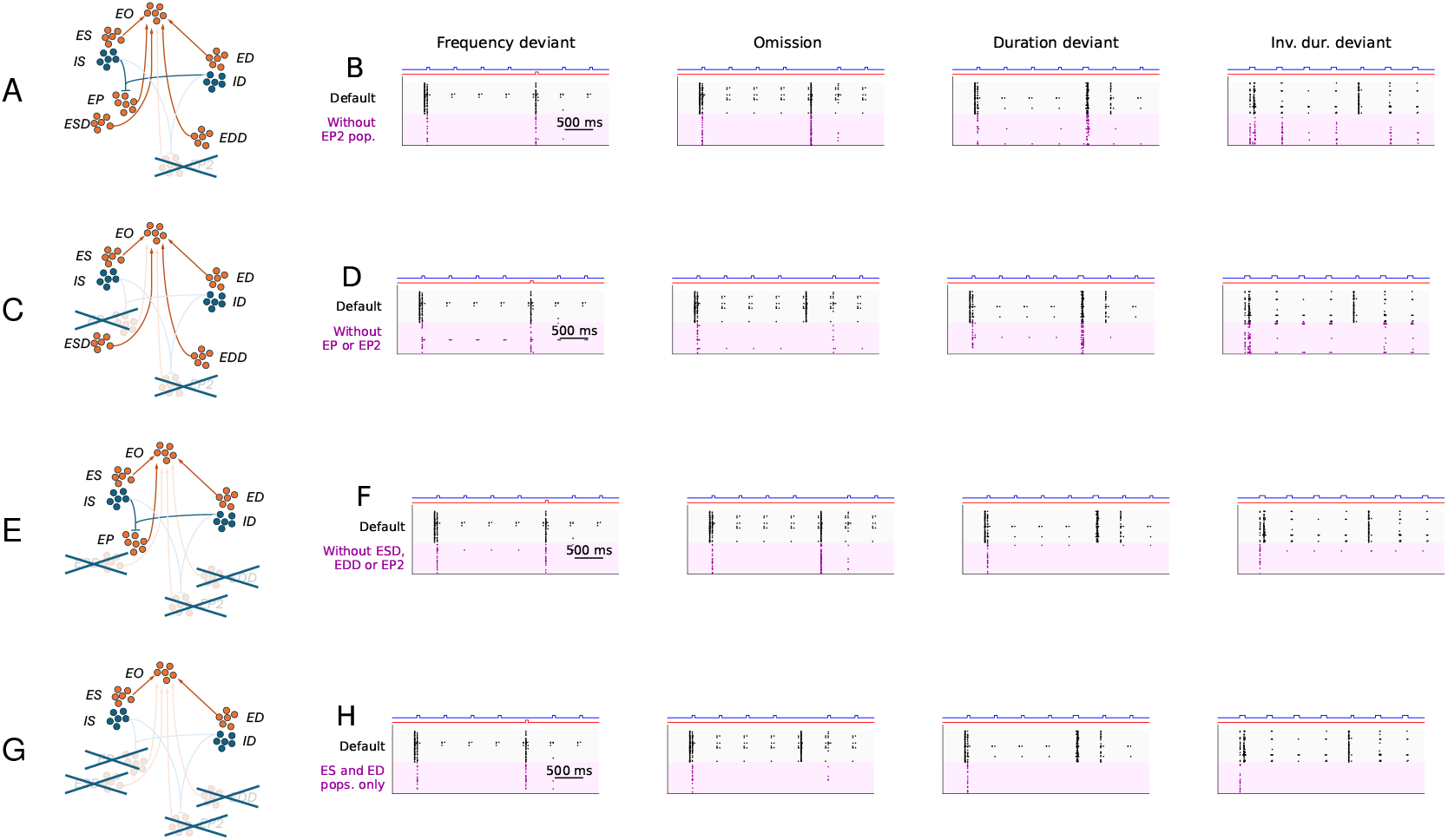
The removal of subpopulations leads to loss of deviance detection in different protocols. **A**,**C**,**E**,**G**: Illustration of the network and the removed populations (crossed out). A: The population EP2 removed. C: The EP and EP2 populations removed. E: The ESD, EDD, and EP2 populations removed. G: The ESD, EDD, EP and EP2 populations removed (this is equal to also IS and ID being removed since they no more interact with any remaining population). **B**,**D**,**F**,**H**: The population spike train of the output population in the default (black) network and in the network where the population(s) indicated in panels A, C, E, and G, respectively, were removed (magenta). The spike trains indicate that the deviance detection was lost in the inverse duration deviant (B), the omission and the inverse duration deviant (D), the duration and inverse duration deviant (F), or the omission, duration and inverse duration deviant (H) protocols.

